# PyMVPD: A toolbox for multivariate pattern dependence

**DOI:** 10.1101/2021.10.12.464157

**Authors:** Mengting Fang, Craig Poskanzer, Stefano Anzellotti

**Affiliations:** Department of Psychology and Neuroscience, Boston College, Boston, MA 02467

**Keywords:** multivariate pattern dependence, connectivity, fMRI, deep networks

## Abstract

Cognitive tasks engage multiple brain regions. Studying how these regions interact is key to understand the neural bases of cognition. Standard approaches to model the interactions between brain regions rely on univariate statistical dependence. However, newly developed methods can capture multivariate dependence. Multivariate Pattern Dependence (MVPD) is a powerful and flexible approach that trains and tests multivariate models of the interactions between brain regions using independent data. In this article, we introduce PyMVPD: an open source toolbox for Multivariate Pattern Dependence. The toolbox includes pre-implemented linear regression models and artificial neural network models of the interactions between regions. It is designed to be easily customizable. We demonstrate example applications of PyMVPD using well-studied seed regions such as the fusiform face area (FFA) and the parahippocampal place area (PPA). Next, we compare the performance of different model architectures. Overall, artificial neural networks outperform linear regression. Importantly, the best performing architecture is region-dependent: MVPD subdivides cortex in distinct, contiguous regions whose interaction with FFA and PPA is best captured by different models.

## 1 Introduction

Cognitive processes recruit multiple brain regions. Understanding which of these regions interact, and what computations are performed by their interactions, remains a fundamental question in cognitive neuroscience. In an effort to answer this question, a large body of literature has used measures of the statistical dependence between functional responses in different brain regions. The most widespread approach – ‘functional connectivity’ – computes the correlation over time between the responses in different brain regions, and has been applied to both resting state fMRI and task-based fMRI [1, 2, 3, 4]. Other methods have been developed to study the direction of interactions, such as Granger Causality [5, 6] and Dynamic Causal Modeling [7].

More recently, the success of multivariate decoding methods [8] and representational similarity analysis [9] has inspired the development of multivariate approaches to study the statistical dependence between brain regions [10]. These approaches take advantage of the rich structure of fMRI response patterns to model the relationships between brain regions. One approach, ‘Informational Connectivity,’ computes the decoding accuracy trial-by-trial for a given categorization in multiple regions and correlates the accuracies across trials [11]. Another method, which can also be applied to resting state studies in which categories for decoding are not available, uses multivariate distance correlation to capture the statistical dependence between regions [12].

Among these novel approaches, MultiVariate Pattern Dependence (MVPD, [13, 14]) is unique in that it trains and tests models of the interactions between brain regions using independent subsets of data, evaluating out-of-sample generalization. Like multivariate distance correlation, MVPD can be applied to both task data and resting state data. Additionally, MVPD can flexibly use a variety of models of dependence, with the potential to incorporate regularization, and to capture linear, as well as nonlinear, interactions between brain regions.

Given the complex nature of the MVPD, a dedicated toolbox can provide researchers with a more accessible entry point to adopt this method. Here, we introduce a freely available open-source toolbox developed in Python: PyMVPD. The toolbox offers an intuitive workflow and a core set of functions for performing MVPD analyses. It also includes Python scripts for a collection of pre-implemented MVPD models and algorithms to evaluate their performance. The algorithms have been designed so that they can be easily customized, enabling users to build their own MVPD models and associated metrics for assessment. With the introduction of the PyMVPD toolbox, we include a novel extension of MVPD using artificial neural networks (implemented in PyTorch), and we provide a comparison of the performance of different models.

The PyMVPD toolbox comes in two versions. The full version is available on Github, and requires a working installation of PyTorch. To use the full version, the use of CUDA and general purpose graphics processing units (GPGPUs) is recommended. In addition, we make available a LITE version of the toolbox, without the neural network models, that does not require PyTorch and can be installed with PyPI.

In the remainder of the article, a brief technical introduction to MVPD is followed by a description of PyMVPD implementation and the analysis workflow (Fig. 1). Next, the algorithms are validated by analyzing a publicly available dataset - the studyforrest dataset [15]. Finally, the performance of different types of models is assessed, taking advantage of MVPD’s flexibility to compare the predictive accuracy of linear regression and artificial neural networks.

**Figure 1:**
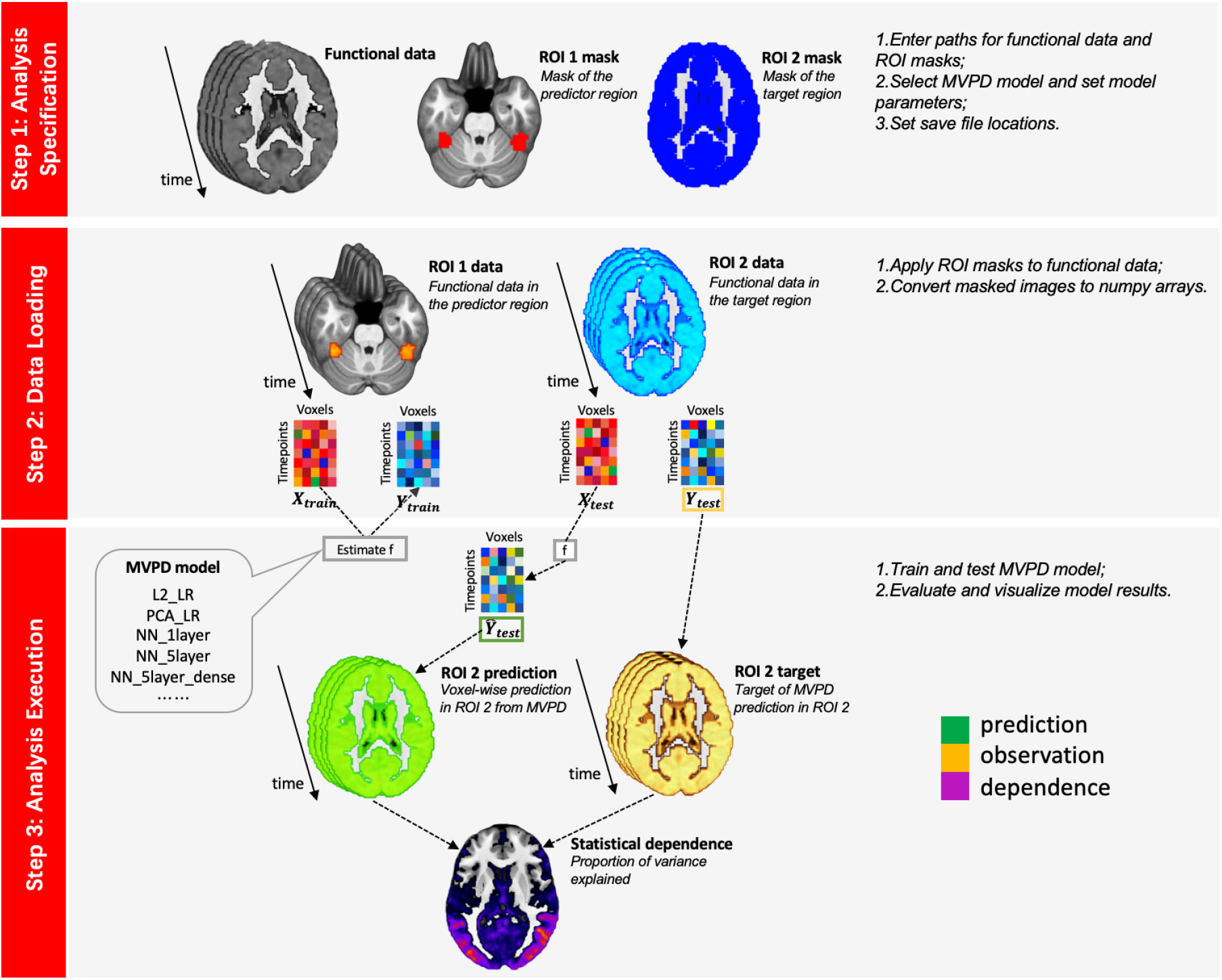
PyMVPD workflow. Analyzing data with the PyMVPD toolbox consists of three steps: (1) analysis specification; (2) data loading; and (3) analysis execution. In step 1, users are required to specify the functional data, masks for both the predictor and the target region, as well as the chosen MVPD model to perform the following analyses. These details can be specified by editing the Python script *run_MVPD.py.* Next, users can proceed with step 2, which loads neuroimaging data and converts it to a suitable format. Finally, step 3 runs the MVPD model and generates the analysis results, which are saved in a user-defined directory.

## 2 Methods

### 2.1 MVPD

Multivariate pattern dependence (MVPD) is a novel technique that analyzes the statistical dependence between brain regions in terms of the multivariate relationship between their patterns of response. Compared with traditional methods used for connectivity analysis, MVPD has two main advantages [13]. First, MVPD preserves the fine-grained information that can be lost by spatially averaging in mean-based univariate connectivity. By doing so, MVPD improves sensitivity as compared to univariate methods such as standard functional connectivity [13]. Second, MVPD is trained and tested with independent subsets of data, therefore it can be more robust to the noise in neuroimaging data.

The logic of MVPD is as follows. Suppose that we wanted to calculate the statistical dependence between two brain regions. MVPD will learn a function that maps the response patterns in one region (the ‘predictor’ region) onto predicted response patterns in the other (the ‘target’ region). Let us consider an fMRI scan with *m* experimental runs. We denote the multivariate timecourses in the predictor region by *X*_1_,…, *X_m_*. Each matrix *X_i_* is of size *n_X_* × *T_i_*, where *n_X_* is the number of voxels in the input region, and *T_i_* is the number of timepoints in the experimental run *i*. Analogously, *Y*_1_,…,*Y_m_* denote the multivariate timecourses in the target region, where each matrix *Y_i_* is of size *n_Y_* × *T_i_*, and *n_Y_* is the number of voxels in the target region.

As a first step, the data is split into a training subset and a test subset. It is important that the training and test subsets are independent. Since fMRI timeseries are characterized by temporal autocorrelation, it is best to not use timepoints from one run for training and adjacent timepoints from the same run for testing. A common approach is to use leave-one-run-out cross validation: this is the approach implemented by default in the PyMVPD toolbox. For each choice of an experimental run *i*, data in the remaining runs is concatenated as the training set

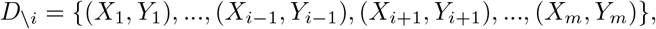

while data *D_i_* = {(*X_i_, Y_i_*)} in the left-out run i is used as the test set.

For convenience, we will denote with *X_train_* and *Y_train_* the concatenated training data in the predictor region and in the target region respectively. The training data is used to learn a function *f* such that

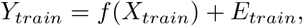

where *E_train_* is the error term. In the current implementation, the response pattern in the target region at a given time is predicted from the response pattern in the predictor region at the same time. However, models that integrate the response in the predictor region across multiple timepoints are a straightforward extension. Once the function *f* has been estimated, we use it to generate predictions of the responses in the target region *Ŷ_test_* given the responses in the predictor region during the test run:

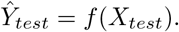

Finally, the accuracy of the prediction is computed. In the PyMVPD toolbox, we provide a measure of predictive accuracy by calculating the voxelwise proportion of variance explained. For each voxel *j* in the target region, variance explained is calculated as:

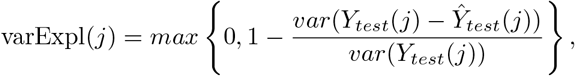

where *Ŷ* are the predicted voxelwise timecourses. The values varExpl(*j*) are then averaged across voxels in the target region and across cross-validation runs to obtain a single measure varExpl. In addition, the PyMVPD toolbox is designed to allow for customized measures of accuracy (more details will be provided in the following sections).

Comparisons between the predictive accuracy of different predictor regions or different types of models can be done using nonparametric statistical tests on the differences between their proportions of variance explained. In the experimental application of PyMVPD in this article, we assessed the statistical significance across participants with statistical nonparametric mapping [16] using the SnPM13 software (http://warwick.ac.uk/snpm). In order to test significance without comparing performance across model types, a control predictor region can be used as baseline for comparison (i.e. a mask for cerebrospinal fluid). Alternatively, phase resampling of the responses in the target of prediction could be used to construct the null distribution (see [17]).

### 2.2 The PyMVPD toolbox

PyMVPD is a Python-based toolbox that implements the MVPD analysis pipeline. This software package is freely available at https://github.com/sccnlab/PyMVPD.git. PyMVPD provides five pre-implemented MVPD models including two linear regression models and three neural network models. The neural network models are built using PyTorch. For users who are only interested in linear regression models, or who would like to avoid the complexities of a PyTorch installation, we have also provided a lite version as PyMVPD_LITE at https://github.com/sccnlab/PyMVPD_LITE.git for which PyTorch is not required. Analyzing data with PyMVPD proceeds in three steps: analysis specification, data loading, and analysis execution.

#### 2.2.1 Step 1 - analysis specification

Prior to MVPD analysis, the fMRI data at hand should have already undergone standard preprocessing steps, such as registration, normalization and denoising. The preprocessed fMRI data should be in NIfTI format. Next, the user should create brain masks of the predictor region (‘RO11’) and the target region (‘ROI2’), also in NIfTI file format.

The MultiVariate Pattern Dependence between brain regions can be computed using a variety of models. Users are required to specify the MVPD model they want to use and to set the corresponding model hyperparameters. The PyMVPD toolbox provides a total of five pre-implemented MVPD models which can be divided into two major categories: linear regression models and neural network models. In addition, users can customize their own models and evaluation metrics by adding scripts to the home directory of the toolbox.

##### 2.2.1.1 Linear regression models

Linear regression attempts to model the relationship between a dependent variable and one or more explanatory variables by fitting a linear function to observed data. Specifically, we view the multivariate timecourses in the predictor region *X* as the explanatory variable and the multivariate timecourses in the target region *Y* as the dependent variable. The MVPD mapping *f* can be modeled with multiple linear regression

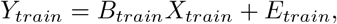

where *B_train_* is the vector of parameters and *E_train_* is the error vector. In PyMVPD, we provide two pre-implemented linear regression models: linear regression with L2 regularization (also known as ridge regression), and linear regression using dimensionality reduced data (with principal component analysis: PCA).

**L2_LR**: A large number of parameters as compared to a relatively small dataset can lead regression models to overfit the data. That is, the model learns a function that corresponds too closely to the particular training set and therefore fails to fit unseen data, resulting in poor predictive accuracy during testing. To mitigate this issue, we provide **L2_LR**, a linear regression model with L2 regularization in the PyMVPD model family.

To implement L2_LR, the linear regression model stays the same, but it is the calculation of the loss function that includes the L2 regularization term:

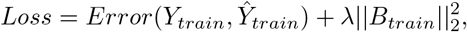

where 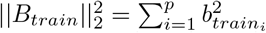 is the square of the L2 norm. This regularization term also introduces a new scalar model parameter λ that adjusts the regularization penalty. The value of λ is set by the user.

**PCA_LR**: Dimensionality reduction provides is an alternative approach to L2 regularization that can be used to mitigate overfitting. Principal component analysis (PCA) is a widely used dimensionality reduction technique that identifies dimensions chosen to preserve as much variance as possible in the original data. The PyMVPD provides **PCA_LR**, a linear regression model that takes as inputs dimensionality-reduced data using PCA. Note that it is also the first version of MVPD implementation [13].

Formally, we first apply PCA to *Y_train_* and *X_train_*:

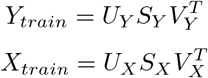

Next, *Y_train_* and *X_train_* are projected on lower dimensional subspaces spanned by the first *k_Y_* and *k_X_* principal components respectively:

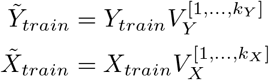

where 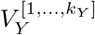 is the matrix formed by the first *k_Y_* columns of *V_Y_* and 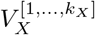 is the matrix formed by the first *k_X_* columns of *V_X_*.

After dimensionality reduction, the mapping *f* from the dimensionality-reduced timecourses 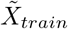 to the dimensionality-reduced timecourses 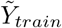 is modeled with multiple linear regression:

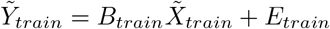

where *B_train_* are the regression parameters. The parameters are estimated using ordinary least squares (OLS).

Once parameters *B_train_* have been estimated, we first projected the test data of the predictor region in the left-out run *i* on the dimensions of the predictor region estimated with the other training runs. To generate the multivariate prediction for the target region in run *i*, we then multiplied the dimensionality reduced test data in the predictor region by the parameters estimated using the other runs during training. More specifically, for each run *i*, we generated dimensionality reduced responses in the predictor region for testing:

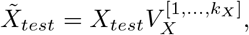

where *V_X_* is calculated using the training data. Then, we calculate the predicted responses in the target region in run *i*:

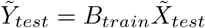

using the parameters *B_train_* independently estimated with the training runs. Finally, we used the inverse of the matrix 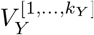 to project the multivariate prediction 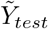 from its low-dimensional subspace of principal components to the original voxel space, obtaining the voxel-wise prediction for the target region:

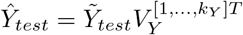

##### 2.2.1.2 Neural network models

In addition to the above linear regression models, we propose an extension in which MVPD between brain regions can be modeled using artificial neural networks. In this approach, the multivariate patterns of response in the predictor region are used as the input of a feed-forward neural network trained to generate the patterns of response in the target region. In PyMVPD, all neural network models are trained using stochastic gradient descent (SGD) on a mean square error (MSE) loss by default. Batch normalization is applied to the inputs of each layer. Additionally, users should set the following hyperparameters for the chosen neural network: number of hidden units, learning rate, weight decay, momentum, minibatch size and epochs for training.

**NN_1layer**: This is the simplest neural network model implemented in the PyMVPD toolbox: a fully-connected linear neural network model with only one hidden layer.

**NN_5layer**: Previous studies have shown that deeper networks can approximate the same classes of functions as shallower networks using fewer parameters [18]. To test the performance of deeper network architectures, PyMVPD includes **NN_5layer**: a pre-implemented neural network model with five fully-connected linear hidden layers.

**NN_5layer_dense**: A challenge encountered in training deep neural networks is the vanishing-gradient problem. Recent architectures designed to mitigate this problem (i.e. DenseNet, [19]) have been extremely effective on a variety of datasets. To test the performance of DenseNet architectures in MVPD, we pre-implemented a five-layer dense neural network model **NN_5layer_dense** with each layer featuring direct connections to all downstream layers.

##### 2.2.1.3 User guide

In the first step of the analysis, the user needs to specify the following details: the participant whose data are to be analyzed (‘sub’); the total number of experimental runs (‘total_run’); the paths to the directories containing processed functional data (‘filepath_func’); the paths to the directories containing the predictor ROI mask (‘filepath_mask1’) and the target ROI mask (‘filepath_mask2’); the path to the directory where the extracted functional data will be saved (‘roidata_save_dir’); the path to the directory where the results will be saved (‘results_save_dir’); the type of model to be used (‘model_type’); the option to save predicted timecourses (‘save_prediction’); and the model hyperparameters.

#### 2.2.2 Step 2 - data loading

The second step of PyMVPD is the loading and processing of input data. Before running the chosen MVPD model, values from the functional data should first be extracted using masks specified in the first step. Then, the extracted functional data should be transformed into numpy arrays in preparation for the following analyses. To accomplish this step, the user simply needs to call the function *load_data()* from *data_loading.py.*

#### 2.2.3 Step 3 - analysis execution

Once the analysis details have been specified and the data is loaded, the third step executes the analysis, estimating the statistical dependence between brain regions and reporting the accuracy of predictions in independent data. To perform step 3, users can call the function *MVPD_exec()* from *model_exec.py*, which will execute the MVPD model and save the results to a folder of their choice (the save folder is specified in step 1).

### 2.3 Application to experimental fMRI data

#### 2.3.1 Data acquisition and preprocessing

As a demonstration of the use of PyMVPD, we analyzed fMRI data of 15 participants (age range 21-39 years, mean 29.4 years, 6 females) watching a movie, from the publicly available *studyforrest* dataset (http://studyforrest.org). Functional data were collected on a whole-body 3 Tesla Philips Achieva dStream MRI scanner equipped with a 32 channel head coil. The high-resolution (3×3×3 mm) BOLD fMRI responses were acquired using a T2*-weighted echo-planar imaging sequence. Complete details can be found in [20].

The dataset includes a movie stimulus session, collected while participants watched the two-hour audio-visual movie ‘Forrest Gump’. The movie was cut into eight segments, and each segment lasted approximately 15 min. All eight segments were presented to participants in chronological order in eight separate functional runs. Additionally, the dataset includes an independent functional localizer that can be used to identify category-selective regions [15]. During the category localizer session, participants viewed 24 unique gray-scale images from each of six stimulus categories: human faces, human bodies without heads, small artifacts, houses, outdoor scenes, and phase scrambled images. Each participant was presented with four block-design runs and a one-back matching task.

All fMRI data was preprocessed using fMRIPrep (https://fmriprep.readthedocs.io/en/latest/index.html). Anatomical images were skull-stripped with ANTs (http://stnava.github.io/ANTs/), and segmented into gray matter, white matter, and cerebrospinal fluid using FSL FAST. Functional images were corrected for head movement with FSL MCFLIRT (https://fsl.fmrib.ox.ac.uk/fsl/fslwiki/MCFLIRT), and were subsequently coregistered to their anatomical scan with FSL FLIRT. Data of one participant was excluded because it could not pass the fMRIPrep processing pipeline. For the remaining 14 participants, we removed noise from the data with CompCor [21] using 5 principal components extracted from the union of cerebrospinal fluid and white matter. Regions of no interest for the cerebrospinal fluid and white matter were defined individually for each participant.

### 2.4 ROI definition

In each individual participant, seed regions of interest (ROIs) in the fusiform face area (FFA) as well as the parahippocampal place area (PPA) were defined using the first block-design run from the functional localizer. We performed whole-brain first level analyses on each participant’s functional data by applying a standard general linear model with FSL FEAT [22]. Next, we identified the peak voxels with the highest t-values for the contrast between the preferred category and other categories (i.e. FFA contrast: faces > bodies, artifacts, scenes, and scrambled images; PPA contrast: scenes > faces, bodies, artifacts, and scrambled images). We generated spheres of 9mm radius centered in the peaks. Finally, the voxels within spheres from the left and right hemisphere were combined, and the 80 voxels with the highest t-values were selected (this is a common choice in neuroimaging studies, see [23, 24].

We additionally created a group-average gray matter mask using the gray matter probability maps generated during preprocessing, with a total of 53539 voxels, that was used as the target of prediction.

### 2.5 PyMVPD analysis

Using the PyMVPD toolbox, we estimated the multivariate pattern dependence between each ROI (FFA/PPA) and the gray matter using all five pre-implemented MVPD models. Specifically, for both FFA and PPA, we took the 80 voxels in the seed ROI as the predictor region, and the 53539 voxels in the gray matter as the target region. For the **L2_LR** model, the regularization parameter λ was set to be 0.001. For the **PCA_LR** model, three principal components were selected. For all the neural network models (**NN_1layer**, **NN_5layer**, **NN_5layer_dense**), we set the number of hidden units in each hidden layer to be 100. Each network was trained with a batch size of 32, a learning rate of 0.001, and a momentum of 0.9 with no weight decay.

For each MVPD model, 7 of the 8 movie runs were used for training, and the remaining run was used for testing. This leave-one-run-out procedure was repeated 8 times by leaving aside each possible choice of the left-out run. We then calculated the variance explained for each voxel in the target region by all five MVPD models in the left-out data.

The proportion of variance explained for each seed region and model was computed for each voxel in gray matter. Next, we compared the relative performance of each pair of MVPD models. For each participant, the proportion of variance explained by each model was averaged across all voxels in gray matter, and across all cross-validation folds. The difference between the average variance explained by the two models was computed for each participant, and significance was assessed with a one-tailed t-test across participants – p-values were Bonferroni corrected for all 20 comparisons (since one-tailed tests were used, comparisons in both directions were counted in the correction).

In addition to testing the model’s overall performance, we sought to determine whether the best-performing models vary depending on the location of the voxels that are the target of prediction. For each voxel in the gray matter, we first selected the model yielding the highest proportion of variance explained in that voxel (averaged across participants) and specified that model as the best model for that voxel. Then, we obtained a conservative measure of the extent to which the model outperformed the other models, calculating the lowest t-value among all comparisons between the best model and all other models.

## 3 Results

### 3.1 PyMVPD with FFA and PPA as seeds

In the first analysis, we tested whether our implementation of PyMVPD can identify expected patterns of statistical dependence between different brain regions. To do this, we selected FFA and PPA as seed regions, and calculated MVPD between these regions and all other voxels in gray matter (Fig. 3). We display the results for a linear 5 layer DenseNet (**NN_5layer_dense**) as model architecture because we expected this architecture to be the most powerful among the pre-implemented models (see Methods for details about the architecture and training procedure); results with other models are similar and are reported in the Supplementary Materials. Using FFA as seed region, we observed statistical dependence (proportion of variance explained > 5% with a broad set of regions including ventral occipital and temporal cortex bilaterally, anterior temporal lobe (ATL), superior temporal sulcus (STS), as well as parietal regions (Fig. 3, top). Despite the fact that we used the FFA in both hemispheres as seed (Figure 2), PyMVPD revealed a right lateralization in statistical dependence in ATL, along the STS, and in parietal regions. Using MNI coordinates extracted from Neurosynth, we established that face-selective regions such as FFA (42 −50 −24; −40 −48 −22), the occipital face area (OFA: 44 −76 −14; −42 −82 −14), the face-selective pSTS (58 −44 12), and the anterior temporal face patch (36 −8 −41; −40 −10 −40) were located within the areas identified by PyMVPD. Within ventral occipito-temporal cortex, statistical dependence with FFA was not restricted to lateral portions of the fusiform gyrus, but extended medially into the parahippocampal gyrus – we discuss this finding in more detail in the Discussion section.

**Figure 2:**
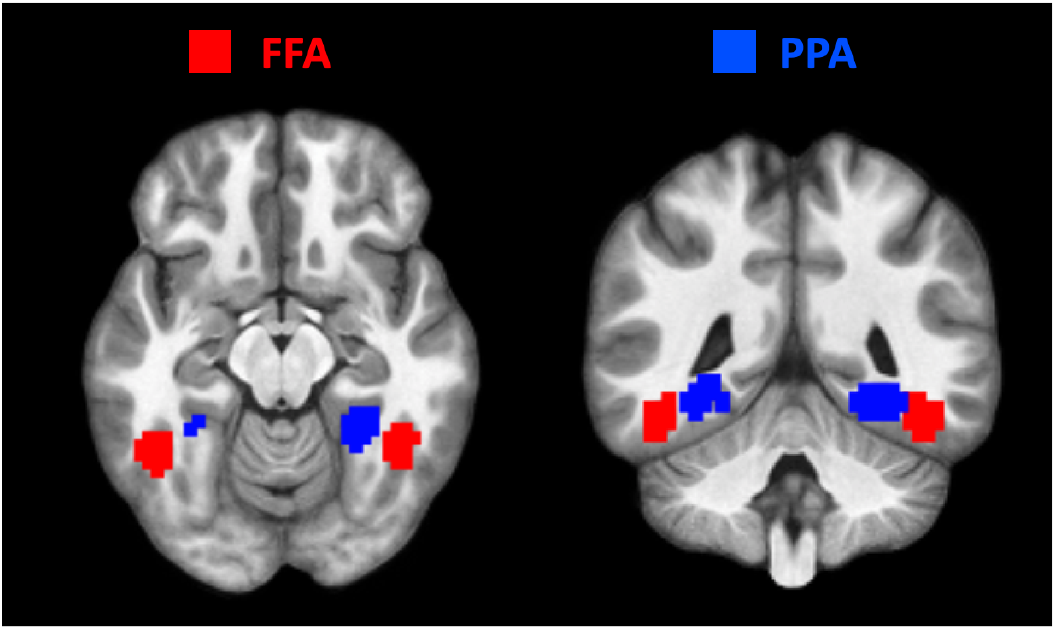
Seed regions. Example seed regions for one representative participant. The fusiform face areas (FFA) are shown in red, and the parahippocampal place areas (PPA) are shown in blue. Individual ROIs per participant were defined based on first-level t-maps, identifying 9mm spheres centered on the peaks for the preferred category, and selecting the top 80 voxels with highest t-values within the spheres.

Using PPA as the predictor region, PyMVPD detected statistical dependence with regions in ventral occipital and temporal cortex, transverse occipital sulcus (TOS), and retrosplenial cortex (Fig. 3, bottom). Using MNI coordinates extracted from Neurosynth, we found that besides PPA (24 −48 −14; −26 −48 −10) which was used as seed, other scene-selective regions such as scene-TOS (32 −84 14; −40 −84 18) and retrosplenial cortex (RSC: 16 −54 16; −16 −54 10) were included among the areas identified. We did not observe evidence of hemispheric lateralization in the set of regions identified by PyMVPD. Within ventral occipito-temporal cortex, statistical dependence was not restricted to the PPA, but extended laterally into the fusiform gyrus (we return to this in the discussion). Overall, the patterns of statistical dependence identified by PyMVPD are in line with previous studies using other methods [25, 26], and reveal stronger interactions between regions that belong to the same category-selective network (i.e. face-selective regions vs scene-selective regions), as well as signs of right-lateralization for face processing.

**Figure 3:**
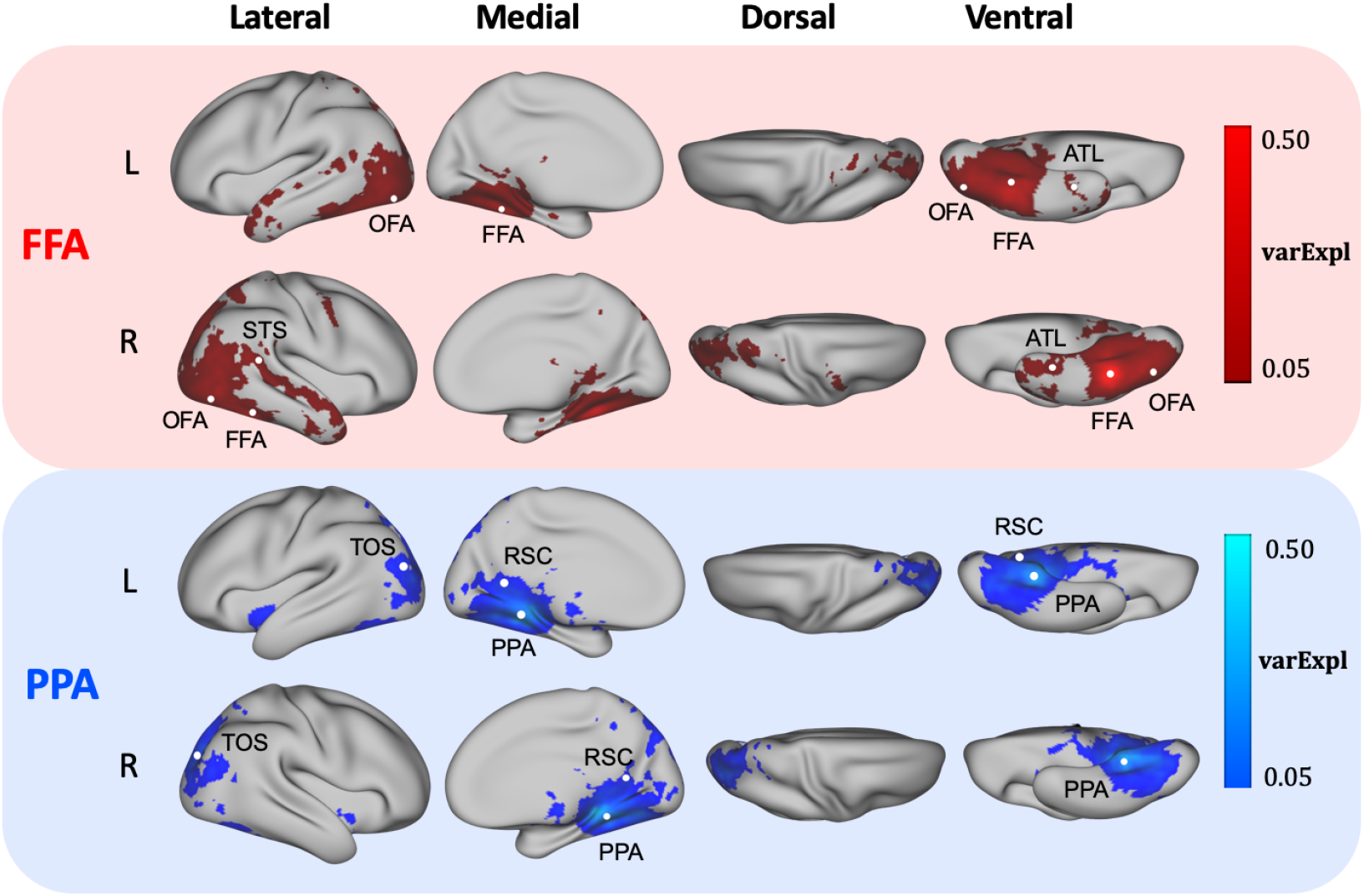
MVPD profiles of the seed regions. Proportion of variance explained varExpl computed by PyMVPD using data from FFA (red) and PPA (blue) as predictors. Results were averaged across all 14 participants. The NN_5layer_dense model was used in this figure. Results from other pre-implemented MVPD models are qualitatively similar and can be found in the Supplementary Materials.

As expected, using FFA as seeds and using PPA as seeds resulted in different whole-brain patterns of statistical dependence: ATL and STS were identified for the FFA seed but not for the PPA seed, and RSC was identified for the PPA seed but not for the FFA seed. In addition to the **NN_5layer_dense** model, other pre-implemented MVPD models in the PyMVPD toolbox also showed similar results (see Supplementary Materials).

### 3.2 Comparing different models

After computing cortical profiles of statistical dependence with FFA and PPA using PyMVPD, we proceeded to compare the predictive accuracy across different MVPD models. The proportion of variance explained for each model was averaged across the whole brain. Then, we performed pairwise comparisons among all five pre-implemented models. For each pair of models, we subtracted the variance explained varExpl of one model from that of another one. This procedure yielded a difference value for each participant, and we conducted a one-sample one-tailed t-test on the difference values across all 14 participants using SnPM. All p-values were Bonferroni corrected for 20 multiple comparisons. Results are shown in Fig. 4 as difference matrices for FFA and PPA respectively.

**Figure 4:**
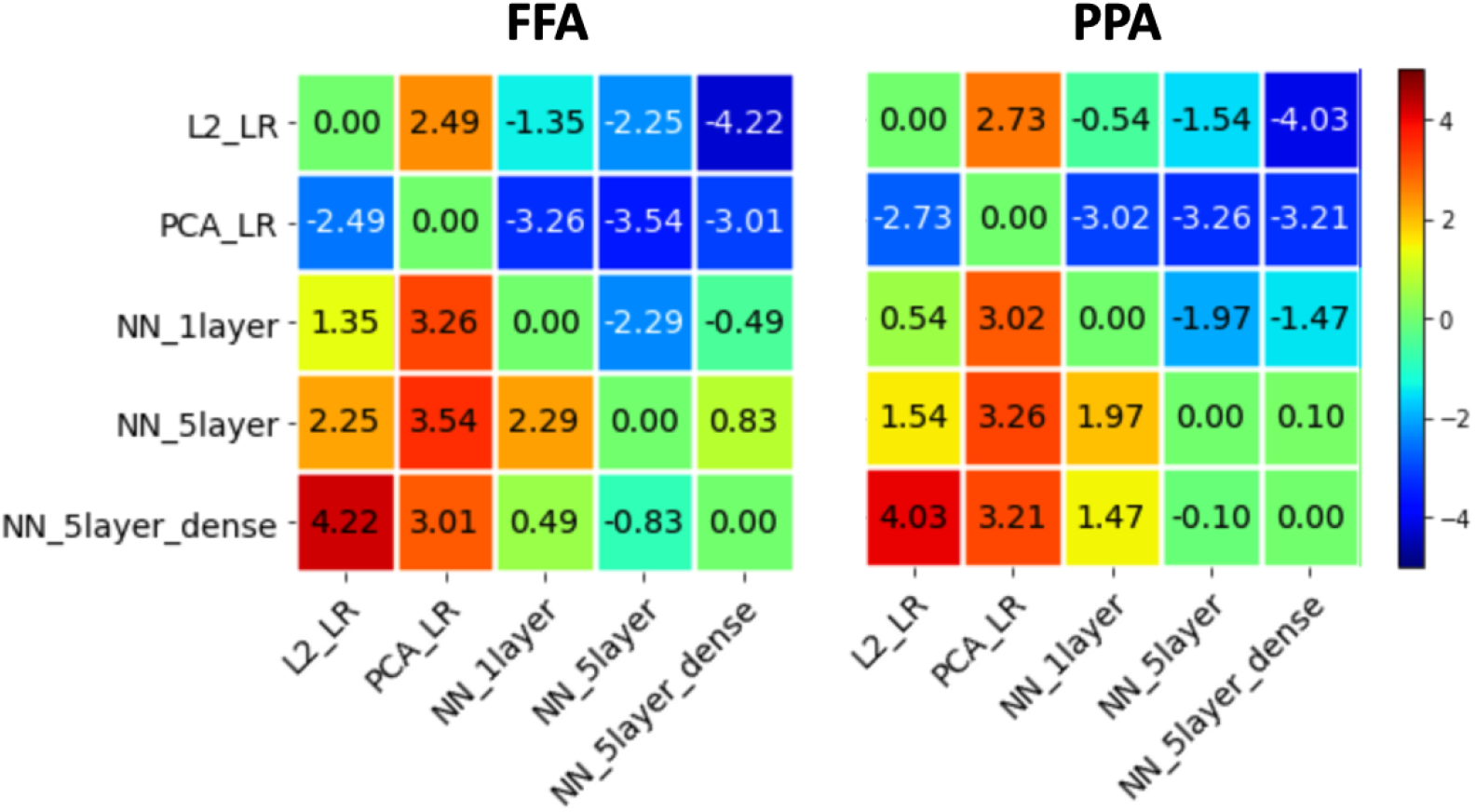
Comparison between models. For each seed predictor region (left: FFA; right: PPA), we plotted the difference matrix of t-values across the five pre-implemented MVPD models. As a measure of overall predictive accuracy, the average proportion of variance explained varExpl was computed across the whole brain for each model per participant. For each pair of models, we subtracted the varExpl of one model from another one, obtaining the pairwise difference values. Finally, we conducted a one-tailed t-test on the difference values across all 14 participants. The corresponding t-values were entered into the difference matrix, indicating the extent to which one model outperformed another one in terms of the overall predictive accuracy using the seed region as the predictor.

Overall, models based on artificial neural networks outperformed standard linear regression models. Linear regression based on principal component analysis (**PCA_LR**) showed the worst predictive accuracy while **NN_5layer** and **NN_5layer_dense** proved to be the two best predicting models. More precisely, using FFA as seed region, **NN_5layer** showed a significantly higher average variance explained than **PCA_LR** (t(13) = 3.54, p = 0.036 corrected) and **NN_1layer** outperformed **PCA_LR** on the borderline of significance (t(13) = 3.26, p = 0.062). **NN_5layer_dense** significantly outperformed **L2_LR** with FFA (t(13) = 4.22, p = 0.010). Using PPA as seed region, **NN_5layer_dense** revealed significantly better predictive performance than **L2_LR** (t(13) = 4.03, p = 0.014). There was also a trend towards significance that **NN_1layer**, **NN_5layer**, and **NN_5layer_dense** outperformed **PCA_LR** (t(13) = 3.02, p = 0.099; t(13) = 3.26, p = 0.062; t(13) = 3.21, p = 0.068) with PPA. The rest of the pairwise comparisons did not show significant differences across participants (p > 0.1).

To further understand the relative accuracy of different models in different brain regions, we generated a map that visualizes the best performing model for each voxel, and the extent to which the best model outperforms the other models (Fig. 5). Specifically, we assigned a different colors to each model (**L2_LR**: green; **PCA_LR**: blue; **NN_1layer**: red; **NN_5layer**: yellow; **NN_5layer_dense**: purple). The color of each voxel was set to the color of the model that performed best at predicting that voxel’s responses, and the color’s saturation was set proportionally to the lowest t-value from all pairwise comparisons between models. In other words, more saturated colors appear in voxels for which the difference between the best model and the runner-up model is greater. This map shows that there isn’t a single best model for all voxels, instead, different voxels are best predited by different models (Fig. 5). **L2_LR** showed as the best-predicting model in the seed regions. **NN_5layer** outperformed other models mainly in the medial and lateral frontal portion of the brain, while **NN_5layer_dense** occupied the ventral and lateral occipital temporal regions.

**Figure 5:**
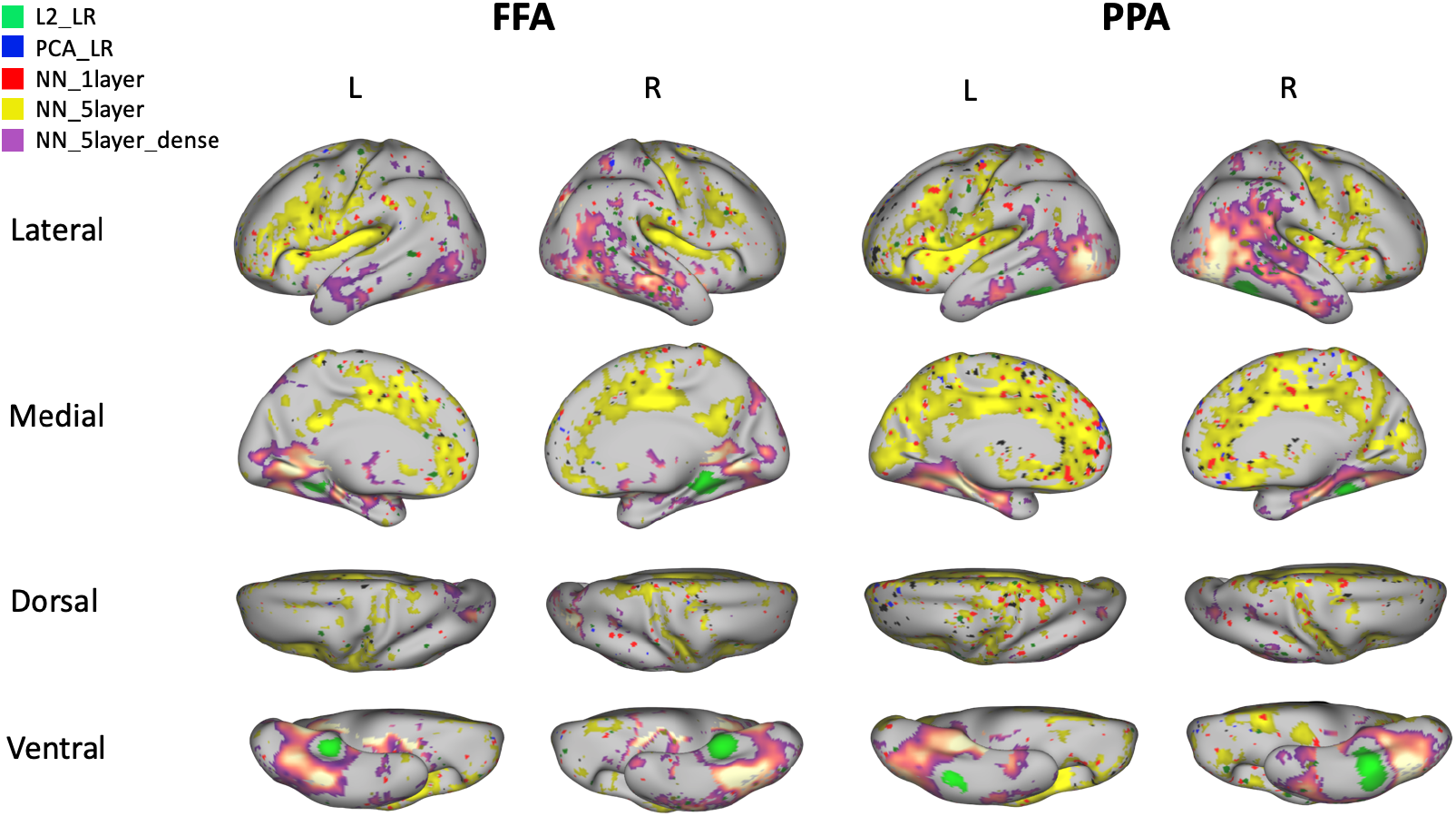
Voxelwise optimal models. For each voxel, the brain maps are colored according to the MVPD model that provides the highest predictive accuracy using seed regions as FFA (left panel) or PPA (right panel) in that voxel. Color saturation reflects the difference between the best model and the second best model. Distinct, contiguous portions of cortex are best predicted by different model types (green: L2_LR, blue: PCA_LR, red: NN_1layer, yellow: NN_5layer, purple: NN_5layer_dense).

## 4 Discussion

In this article, we have introduced PyMVPD, a Python-based toolbox for Multivariate Pattern Dependence (MVPD). MVPD is a novel technique that investigates the statistical relationship between the responses in different brain regions in terms of their multivariate patterns of response [13]. Previous studies have shown that this approach brings higher sensitivity in detecting statistical dependence than standard functional connectivity [13, 27]. However, given the complex nature of the analysis, the implementation of MVPD can be an obstacle. The PyMVPD toolbox makes MVPD more accessible to a broader community of researchers.

PyMVPD provides users with a flexible analysis framework to study the multivariate statistical dependence between brain regions, a collection of pre-implemented MVPD models, and associated metrics to evaluate the analysis results. As a user-friendly toolbox, PyMVPD enables researchers to perform complex MVPD analyses with a few lines of easily readable Python code.

One key property of PyMVPD is that it permits the use of a variety of models to study the multivariate statistical dependence between brain regions. In addition to the standard linear regression models that have proven to be efficient in previous literature [13], we make artificial neural networks available for connectivity research through an integration with PyTorch. The artificial neural network version of MVPD implemented in PyTorch is introduced for the first time in this article. We demonstrate that the neural network implementations of MVPD outperform the previously published version based on PCA in most brain regions. Three neural network architectures come already implemented in PyMVPD, and the toolbox is designed so that it is straightforward to program new custom architectures and integrate them with the other scripts.

In the experimental applications described in this work, we tested PyMVPD using the studyforrest dataset, with the FFA and PPA as seed regions. The results revealed interactions between these seed regions and the rest of the brain during movie-watching, following a pattern that is consistent with the previous literature. Category selective peaks identified with Neurosynth fell within the MVPD maps for the corresponding category. Overall, artificial neural networks outperformed linear regression models in terms of the predictive accuracy for statistical dependence. Importantly, this is not a trivial consequence of the fact that the artificial neural networks are more complex. In fact, MVPD trains and tests models with independent subsets of the data, and models with more parameters do not necessarily perform better at out-of-sample generalization.

Interestingly, no single model that outperformed all others in every voxel. In particular, the **NN_5layer** identified the greatest statistical dependence between seed regions and medial frontal and parietal regions, as well as lateral frontal regions and the insula. By contrast, the responses in the ventral and lateral occipito-temporal regions were best explained by **NN_5layer_dense**.

Taken together, these results indicate that the statistical dependence between different sets of regions might be best predicted by different models. In the literature on brain connectivity, to date, the focus has been on whether or not two brain regions interact. However, a key direction for future research consists in investigating not only whether two regions interact, but also how they interact. The observation that the statistical dependence between the seed regions and different voxels were best captured by different models suggests that PyMVPD could be used to make progress in this direction: studying what model provides the best account for the interaction between two brain regions could yield insights into the transformations that relate the information they encode.

To pursue goals such as this, PyMVPD is designed to be easily customized and extended. In addition to the five pre-implemented models (i.e. **L2_LR**, **PCA_LR**, **NN_1layer**, **NN_5layer**, **NN_5layer_dense**), PyMVPD allows users to customize their own MVPD models as well as evaluation metrics, making this toolbox an ideal environment to compare the predictive accuracy of different types of models to study the interactions between brain regions.

Installing the full version of PyMVPD requires a working installation of PyTorch, installed compatibly with the version of the CUDA drivers of the GPUs. For users who prefer to avoid this step and do not need to use the neural networks, we make available the LITE version of PyMVPD, that includes only the linear regression models, and does not require PyTorch. The LITE version can be also installed using the Python Package Index (with ‘pip’).

Together with both versions of PyMVPD, we provide tutorials on how to calculate MVPD using the toolbox (https://github.com/sccnlab/PyMVPD/blob/main/exp/PyMVPD_Tutorial.ipynb, https://github.com/sccnlab/PyMVPD_LITE/blob/main/exp/PyMVPD_LITE_Tutorial.ipynb). The tutorials are written with Jupyter Notebook, and include sample data as well as the option to plot one’s results side by side with the result we computed. This will make it easier for users to check that the toolbox was installed correctly and to confirm that the results match with those we obtained.

The present study focused on the FFA and PPA as seed regions because they have been studied in depth in previous literature. Future studies can extend our results, investigating the application of PyMVPD to other seed regions. The current implementation of PyMVPD is based on simultaneous prediction: responses in the target region at a given time are predicted from responses in the predictor region at the same time. However, other researchers could take advantage of the customization options to use the responses in multiple timepoints in the predictor region to predict the responses in the target region at each timepoint. Finally, the models of statistical dependence implemented by PyMVPD are deterministic. Multivariate probabilistic models that capture the distribution of uncertainty in predictions are in principle possible, but would require large amounts of data for training.

Although PyMVPD was specifically developed for fMRI analysis, the generic design of the framework makes it widely applicable to other data acquisition modalities (i.e. EEG, MEG) across a variety of domains of brain imaging research. We hope that this toolbox removes some of the barriers to the adoption of MVPD, and facilitates the diffusion of multivariate analyses of the interactions between brain regions.

### Information Sharing Statement

The source code of the presented Python-based toolbox, PyMVPD, is freely available for non-commercial use from https://github.com/sccnlab/PyMVPD.git. The data that support the experimental applications of PyMVPD are available in the publicly released studyforrest dataset at http://studyforrest.org.

## Acknowledgments

We would like to thank Aidas Aglinskas, Emily Schwartz, and Tony Chen for their comments on a previous version of the manuscript, and the researchers who contributed to the *studyforrest* project (Hanke et al., 2016; Sengupta et al., 2016) for sharing their data, and the developers of fmriprep (Esteban et al., 2018) for their assistance with the fmriprep preprocessing pipeline. We would also like to thank Kyle Kurkela for his contributions on the improvement of the code. This work was supported by a startup grant from Boston College and by NSF grant 5109521 to Stefano Anzellotti.

## Supplementary Materials

### Supplementary Figures

**Figure S1:**
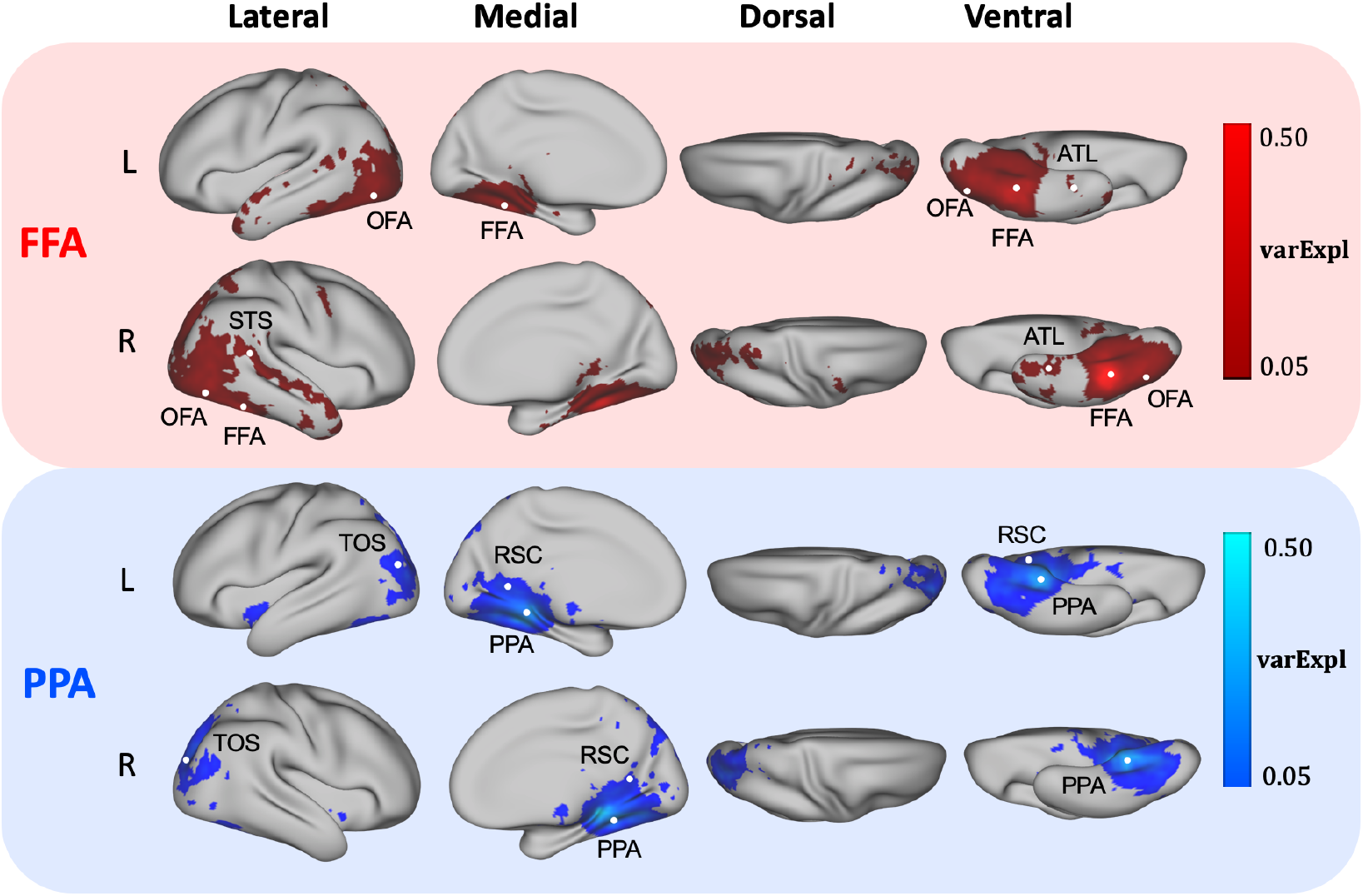
Predictive accuracy of the **L2_LR** model. The proportion of variance explained varExpl obtained by PyMVPD using data from FFA (red) and PPA (blue) as predictors was averaged across all 14 participants and was thresholded at 5%.

**Figure S2:**
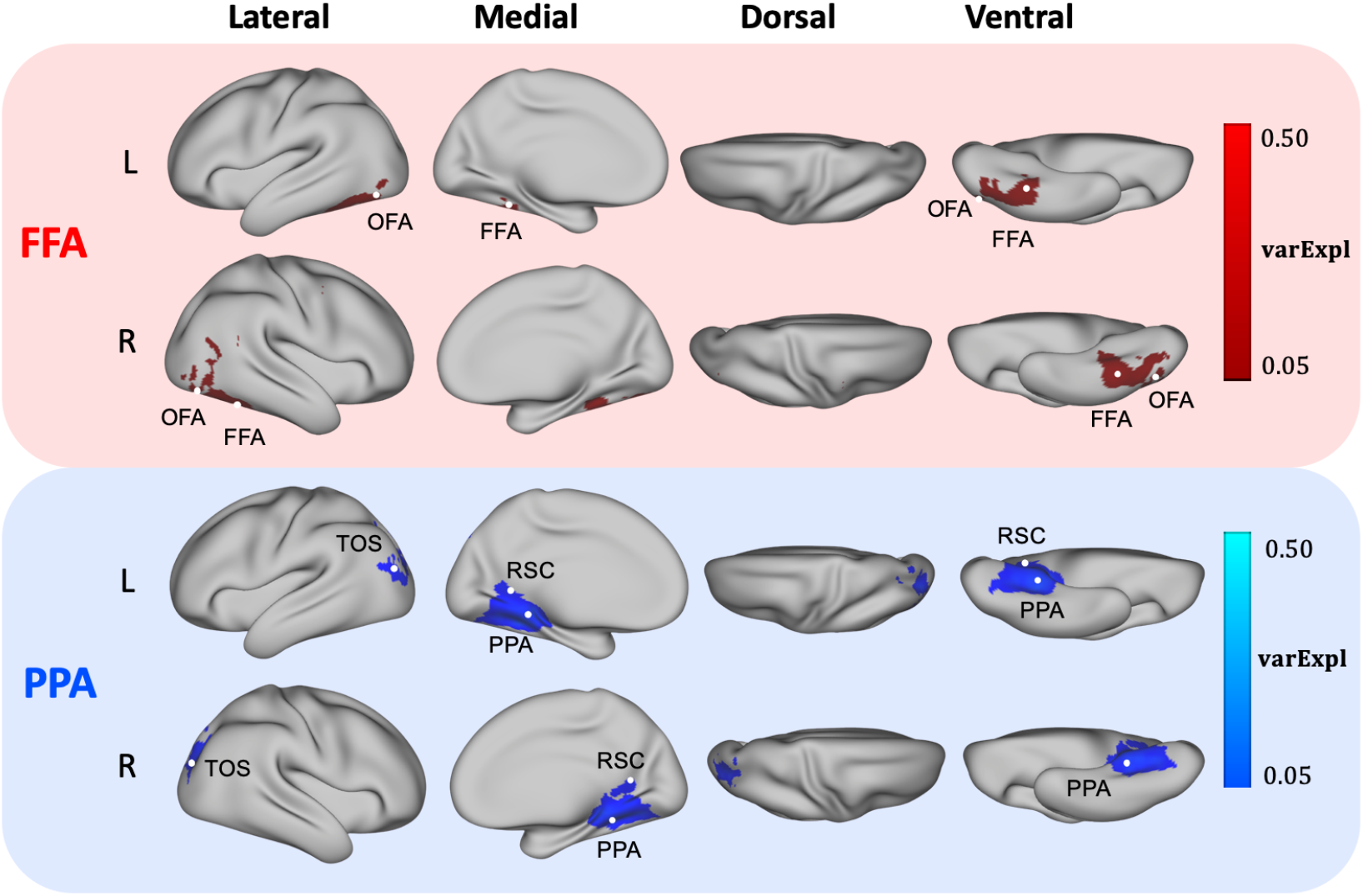
Predictive accuracy of the **PCA_LR** model. The proportion of variance explained varExpl obtained by PyMVPD using data from FFA (red) and PPA (blue) as predictors was averaged across all 14 participants and was thresholded at 5%.

**Figure S3:**
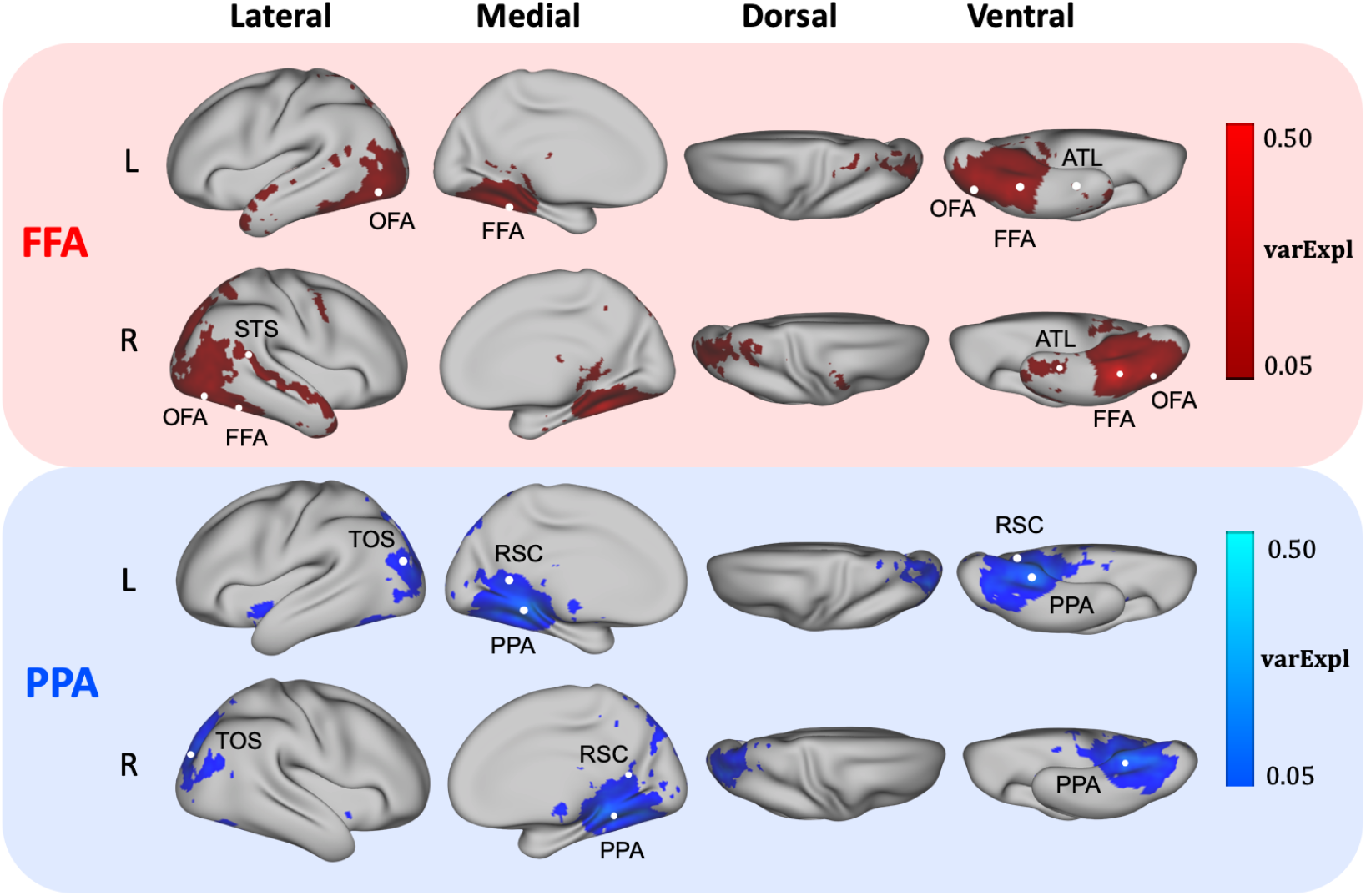
Predictive accuracy of the **NN_1layer** model. The proportion of variance explained varExpl obtained by PyMVPD using data from FFA (red) and PPA (blue) as predictors was averaged across all 14 participants and was thresholded at 5%.

**Figure S4:**
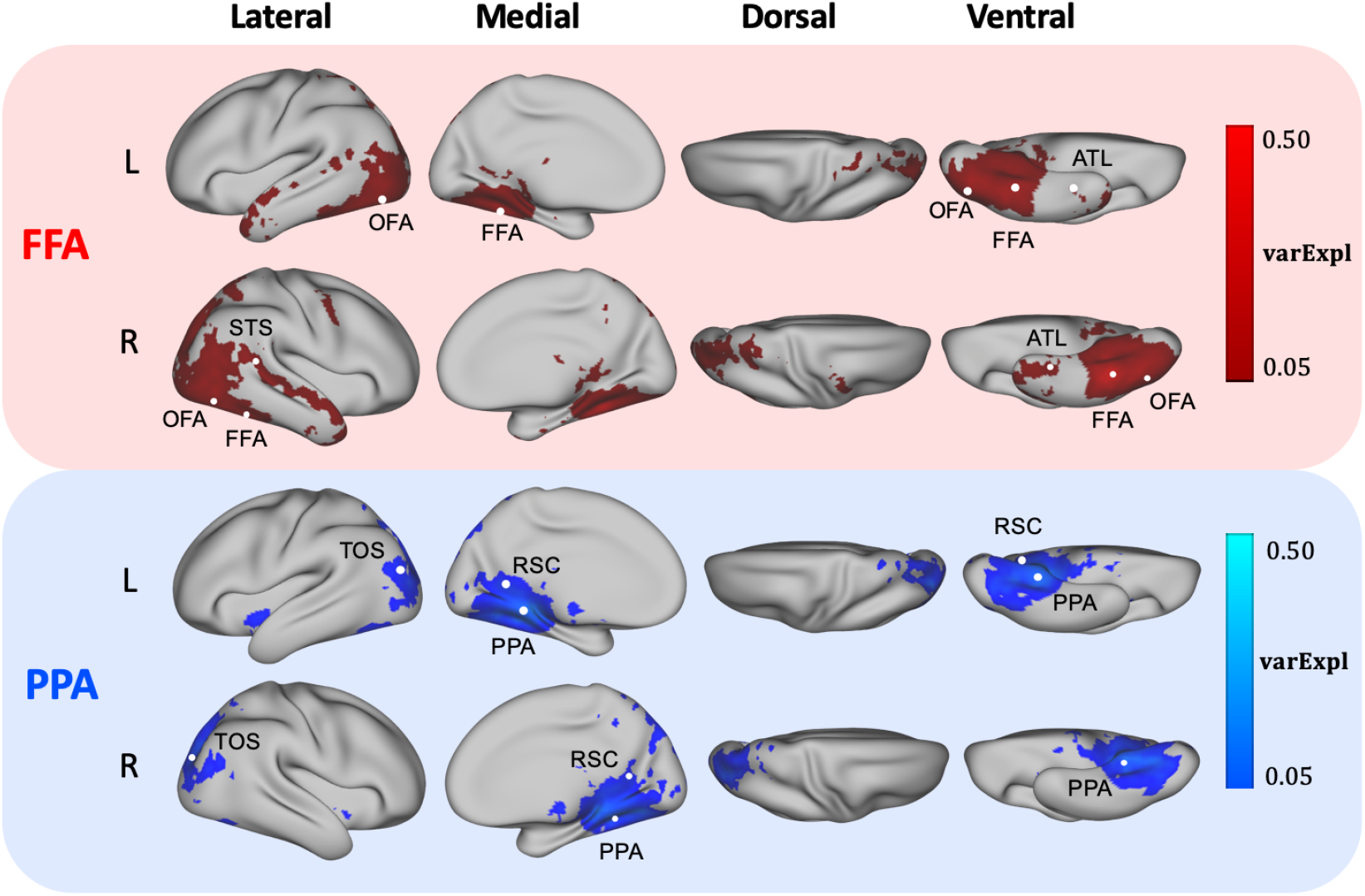
Predictive accuracy of the **NN_5layer** model. The proportion of variance explained varExpl obtained by PyMVPD using data from FFA (red) and PPA (blue) as predictors was averaged across all 14 participants and was thresholded at 5%.

## Notes

### Competing Interest Statement

The authors have declared no competing interest.

